# Deciphering the microbial landscape of lower respiratory tract in health and respiratory diseases

**DOI:** 10.1101/2025.06.11.659233

**Authors:** Cheng Cheng, Yangqian Li, Suyan Wang, Haoyu Wang, Dan Liu, QingLan Wang, You Che, Linlin Xue, Ningning Chao, Xuan He, Chengping Li, Huohua Tian, Jing Zhou, Xin Wang, Yurui Yang, Yuqi Zhu, Renjie Xu, Zhoufeng Wang, Weimin Li

## Abstract

The microbiota plays an important role in maintaining lung homeostasis and the development of respiratory diseases. However, there have not been detailed characteristics of lung microbiome in health and respiratory diseases. Bronchoalveolar lavage fluid of 43 healthy individuals, 23 COVID patients, and 28 non-small cell lung cancer patients was conducted to characterize microbial diversity and function by metagenomic sequencing. We identified 196 species in health lung after removing low abnudance species. The most abundant species were *Staphylococcus aureus*, *Salmonella enterica*, and *Klebsiella pneumoniae*. Keystone species were identified through co-network analysis, such as *Granulicatella adiacens* and *Mogibacterium diversum*. To further explore the microbial function in the healthy lung, we obtained 56 non-redundant metagenome-assembled genomes (MAGs) and subsequently classified them into nine phyla. Metabolism and genetic information processing exhibit high abundance, including bacterial secretion systems, protein export, and *Staphylococcus aureus* infection, indicating a close correlation between microbiota and the host. Furthermore, the microbiome pairwise distances of non-small cell lung cancer (NSCLC) participants were more similar to those of the healthy group than those of individuals with COVID-19. *Rothia dentocariosa*, *Corynebacterium kefirresidentii, Sphingobium yanoikuyae* were enriched in the healthy group, while *Schaalia odontolytica, Actinomyces graevenitzii,* and *Rothia mucilaginosa* was enriched in the NSCLC group. Overall, this study provides a comprehensive overview of the diversity and gene function of the lung microbiome, which will facilitate future studies of microbiota associated with respiratory diseases in humans.

## Introduction

The lung microbiota plays an important role in the human lung respiratory system[1]. Numerous studies have indicated that the diversity and composition of the respiratory microbiome in healthy individuals exhibit individual variability but are generally stable [2–6]. Using high-throughput sequencing, scientists can more accurately describe the composition of the lung microbiota in healthy individuals, including the abundance and functions of microorganisms [7]. Furthermore, the development of metagenomic binning technology has enabled the acquisition of nearly complete metagenome-assembled genomes (MAGs) on a large scale [8]. The microbiota composition in healthy human lungs has been partially characterized primarily through 16SrRNA amplicon sequencing [9, 10]. However, there have been limited assemblies of MAGs specifically targeting the healthy lung microbiome.

Due to clinical sample acquisition difficulty in healthy individuals and limited sequencing depth, the availability of sequenced genomes and functional information for most healthy individual lung microbes remains limited. In our previous studies, we showed pulmonary microbiome dysbiosis in lung cancer patients based on non-assembled metagenomic data[11, 12]. A large-scale and high-depth metagenomic sequencing scan is needed to perform to characterize microbiota compositions and functions in healthy human lungs. Here we address this problem by constructing co-network of bacterium based on their relative abundance, reconstructing draft prokaryotic genomes from 43 healthy individuals. Furthermore, we revealed the microbiota alterations in lung diseases focused on COVID-19 and non-small cell lung cancer (NSCLC) using microbial profiling of this healthy cohort. Our study will provide more comprehensive baseline data for research on human pulmonary microbiota, and better understand its mechanisms of action in both health and disease states.

## Methods

### Sample collection and processing

All details about the experimental samples are listed in Table S1, including 43 healthy individuals, 23 COVID-19 patients, and 28 NSCLC patients. Healthy individuals were recruited from Suining Central Hospital, Sichuan Province, in 2021. They underwent CT imaging, pulmonary function tests, electrocardiograms, and routine blood tests, all showing normal liver and kidney function.

COVID-19 patients were diagnosed with SARS-CoV-2 infection through qPCR or antigen testing. NSCLC patients were histologically and endoscopically confirmed by examination from three pathologists. The samples of those patients were obtained from the West China Hospital of Sichuan University. All patients were followed up to confirm the final diagnosis.

In this study, samples were collected via bronchoscopy for distal alveolar lavage. We used only bronchoscope working channel washes, which were done twice with a minimum volume of 15 mL of 0.9% saline solution. Immediately dispense the collected BALF into 1.8 mL sterile freezing tubes and store at –80°C until processing. The collection process was carried out following aseptic procedures to prevent contamination from environmental, human commensal, and miscellaneous bacteria.

### DNA extraction and quality control

DNA extraction from BALF was performed using the DNA Microbiome kit (51704, Qiagen, USA) according to the manufacturer’s instructions. The host cells are differentially lysed, followed by enzymatic digestion, to efficiently remove host DNA. Subsequently, complete cell lysis is achieved through physical and chemical methods, facilitating the isolation of bacterial DNA.

### Library construction and sequencing

Using an NGS Nextera™ DNA Flex Library Prep kit (Illumina), the DNA libraries were conducted. By performing an enzymatic reaction called segmentation, DNA is fragmented, followed by the addition of adapter sequences and PCR amplification. An Agilent 4200 Bioanalyzer (Agilent Technologies, Santa Clara, USA) was used to assess the quality of the DNA libraries. After a quality check, Illumina PE150 sequencing was conducted by pooling multiple libraries based on the required effective concentration and target data amount. Sequencing was performed on the Illumina NovaSeq 6000.

### Data quality control

For the quality control of raw data obtained from the Illumina sequencing platform, mocat2 (v2.1.3) was used to filter out low-quality reads, and reads aligned to the GRCh38 Homo sapiens reference genome were removed [34]. The remaining high-quality reads were further analyzed.

### Taxonomic classification

Taxonomic classification was conducted with Kraken2 using the standard Kraken2 database (https://benlangmead.github.io/aws-indexes/k2) containing archaea, bacteria, viruses, plasmids, human1, and UniVec_Core. Bracken [35] was then used to calculate the abundance of the identified species from the Kraken2 analysis.

### Diversity analysis

Bacterial diversity was assessed by calculating the Shannon index using the vegan package (v2.5.7) in R. The Wilcoxon test was conducted to determine the statistical significance of differences among groups of healthy individuals. Differences associated with a P-value of < 0.01 were considered significant.

### Co-occurrence network construction

Spearman’s correlation coefficients of relationships across species with an absolute value > 0.8 and statistically significant (P-value < 0.01) were visualized using the R package igraph (v2.0.2) with a layout based on the Fruchterman-Reingold algorithm. We also calculated degree and page rank values to facilitate the identification of keystone species. Network module detection was performed using the greedy algorithm.

### Microbiome-functional pathway analysis

To identify microbial pathways, the human removed reads were analysed with HUMAnN (v3.7) [36] by using the databases of uniref 90[37] and pathway MetaCyc[38]. The “WGCNA” package[39] was used to conduct WGCNA (weighted gene co-expression network analysis). For correlation analysis, spearman correlations were calculated using function corr.test in R.

### Metagenomic de novo assembly and binning

The Megahit module from metaWRAP-Binning was utilized to assemble clean data from the aforementioned processed Illumina sequencing data [40]. Metagenomic binning was subsequently conducted using MaxBin2, metaBAT2, and CONCOCT [40]. DAS Tool (v1.1.5) was used to aggregate multiple binning predictions into a new and enhanced bin set [41]. CheckM2 was used to evaluate the quality of the assembled bins, which were screened based on the criteria of completeness (>50%) and contamination (<10%)[42]. Then dRep (v3.0.0) was then used to remove duplicate bins with the parameter “-sa 0.95”. After analysis, 56 MAGs were retained.

### Taxonomic assignment and phylogenetic analysis

MAGs were taxonomically classified using the Genome Taxonomy Database Toolkit (GTDB-Tk) (v2) with “classify_wf” function and default parameters[43]. The Average Nucleotide Identity (ANI) values between the five new MAGs and genomes from two reference cohorts were calculated using fastANI (v1.33). All phylogenetic trees of the 56 MAGs were constructed using iqtree2 (v2.2.3) (https://github.com/iqtree/iqtree2). Finally, iTOL (https://itol.embl.de) was used to generate a phylogenetic tree of the 56 MAGs.

### Function annotation of MAGs

High-quality MAGs were utilized for gene prediction by Prodigal (v2.6.3) software (https://github.com/hyattpd/Prodigal) [44]. All complete genes were clustered at 90% protein identification using CD-HIT (v4.8.1) [45]. The KEGG (Kyoto Encyclopedia of Genes and Genomes) annotation results were extracted using KofamKOALA (v1.3.0) software [46]. Additionally, key virulence factors were identified using the Virulence Factor Database (VFDB) [47] via BLAST (v2.10.1+).

### Calculation of the abundances of MAGs and genes

The quant_bins module from metaWRAP-Binning was used to quantify the abundance of each MAG in each sample. BWA MEM was used to align clean reads from each sample to gene catalogs after removing human contamination [48]. The abundances were normalized to fragments per kilobase of gene sequence per million reads mapped (FPKM). The abundances of microbial taxa, KEGG pathways, and Virulence Factors (VFs) were calculated by combining the abundances of all the members within each category.

### Microbiome analysis of three groups

Principal Coordinates Analysis (PCoA) based on the Bray–Curtis distance matrix was utilized to visualize the compositional profiles of 212 species from samples belonging to three groups. The differences in microbiome composition between different phenotypes were calculated using permutational multivariate analysis of variance with distance matrices in the adonis2 function of the vegan package, with 999 permutations.

Associations of specific microbial species with phenotypes were calculated using multivariate analysis by linear models (MaAsLin2) (http://huttenhower.sph.harvard.edu/galaxy) with healthy and lung-diseased samples as references. The p-value of associations was computed by MaAsLin2, and the false discovery rate (FDR) was calculated using the Benjamini–Hochberg correction. FDR < 0.01 was considered significant. Furthermore, the fold change was calculated using R (v4.3.2) with the Wilcoxon test.

## Results

### Distribution of species composition in the healthy human population

We conducted meta-genomics profiling of bronchoalveolar lavage fluid (BALF) from 43 healthy individuals, including 11 males with smoking history (Figure 1a-b; Table S2). We obtained an average of 4,683,836,943 bases in each sample after quality control and removal of human-derived reads (Table S2). Using Kraken2 profiling, we identified 196 species with a relative abundance higher than 0.01 and present in more than 5% of individuals, accounting for 0.991 of the relative abundance per sample from healthy individuals (Table S3-4). Those 196 species were assigned to eight phyla, with the dominant phyla being *Firmicutes*, *Proteobacteria*, and *Actinobacteria* (Figure 1c). In Ibironke et al.’s study, the three phyla mentioned above were identified as dominant phyla in the human respiratory microbiome of five individuals based on full-length 16SrRNA sequencing[13]. To identify microbial species may play a critical role in the maintaining of lung homeostasis, we examined our cohort for microbial taxa with high abundance and core microbes present in over 95% of individuals. We detected 15 high-abundance species as dominant species (Figure 1d-e). Meanwhile, we observed 32 core species (>5% of individuals) out of 196 species. 12 out of the 32 core microbes are also dominant species, indicating their critical roles in the lung ecosystem.

**Figure 1.**
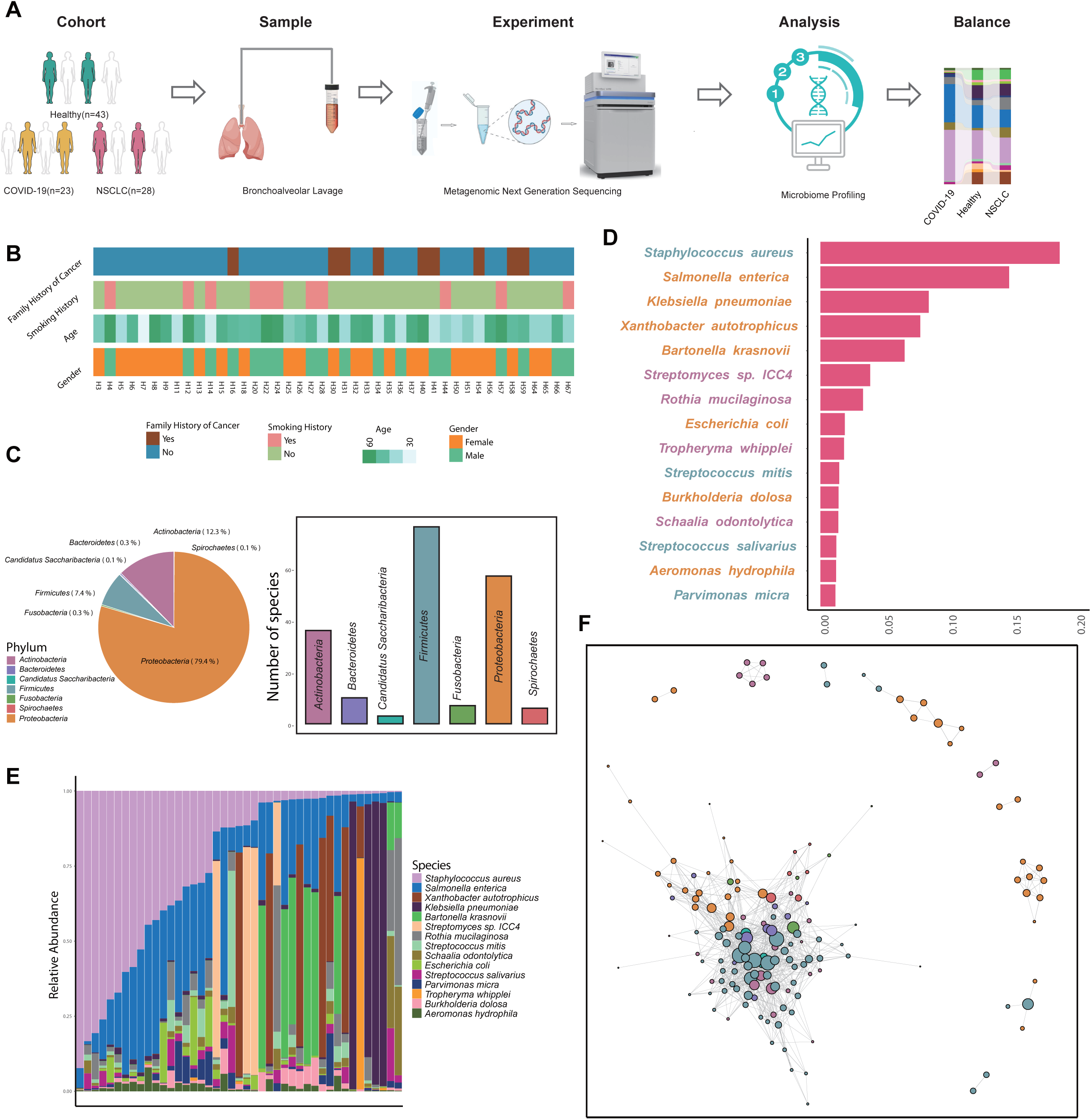
Overview of the microbiome composition of healthy individuals. A. A schematic representation of the workflow of metagenomic sequencing on healthy individuals. B. The clinical information of 43 BALF samples from healthy volunteers including gender, age, history of smoking, and history of family-member cancer. C. (1) Average relative abundances of bacterial phyla after filtering out rare species (those with a relative abundance lower than 0.01 or present in less than 5% of individuals). (2) Number of species present in each phylum. D. Top 15 species-level abundances of all samples in the cohort are sorted by the mean relative abundance of bacteria at the species level in each person. The scale of 0–1 corresponds to 0– 100% abundance. The color of the species name indicates the phyla, as shown in Figure 1c legend. E. Top 15 species-level compositions of all samples in the cohort, sorted by the abundance of *Staphylococcus aureus*. Each vertical line indicates one sample. F. Co-occurrence network of 155 species based on Spearman correlation coefficients among those species. Each node represents one species. Ecological relationships are represented by edges connecting two nodes. Grey lines indicate a positive correlation. The node color represent phyla and the node size is proportional to the rank value.

We further investigated the microbial component in our cohort through principal coordinate analysis (PCoA) and founded that there was no significantly separation based on gender, age, smoking history, or family cancer history (Figure S1a). Furthermore, the alpha diversity of individuals grouped by the aforementioned clinical features was insignificant, indicating that differences in microbial communities between clinical groups were minor (Figure S1b).

To identify keystone species and explore relationships among species. Spearman’s correlation coefficients were calculated, and the ecological relationships across 155 different microbes (a relative abundance > 0.001 and present in more than 60% of individuals) were visualized using co-occurrence networks. There were 155 nodes and 1084 edges (Figure 1f). The top five species with the highest degree and page rank values were *Granulicatella adiacens*, *Mogibacterium diversum*, *Streptococcus australis*, *Mogibacterium pumilum*, and *Streptococcus sp.116-D4* (Figure 1f). Two dominant species, *Staphylococcus aureus* and *Salmonella enterica*, have relatively low degree and page rank values (Figure S1d). Additionally, 12 communities were detected (Figure S1c). *Firmicutes*, *Proteobacteria*, and *Actinobacteria* comprised the three largest communities. Module one was dominated by *Proteobacteria*, module two by *Bacteroidetes* and *Firmicutes*, and module three by *Firmicutes* and *Actinobacteria*. The phylum of the species was significantly associated with the corresponding module (P value < 0.001, Fisher test), suggesting that species within the same phylum may perform similar roles in the healthy lung micro-ecology. The results indicate that the three major phyla (*Firmicutes*, *Proteobacteria*, and *Actinobacteria*), based on abundance and co-network analysis played a crucial role in lung homeostasis.

### Metabolic profiling of microbe from the healthy human lung

To illustrate the metabolic composition of healthy individuals, we identified 374 pathways from the MetaCyc database. The top 15 relative abundances of metabolic terms are shown in Figure 2a-b. These dominant metabolic terms include aerobic respiration I, aminoimidazole ribonucleotide biosynthesis, adenosine nucleotides de novo biosynthesis, guanosine ribonucleotides biosynthesis, and others. We conducted weighted gene co-expression network analysis (WGCNA) to explore the relationship between the 374 pathways and basic clinical information. Two modules were obtained (Figure S2a). However, we did not find modules significantly related to the basic clinical information (Figure S2b). Ethanolamine Utilization and Pyruvate Fermentation to Isobutanol was enriched in the female group compared to the male group (Figure 2c). UMP biosynthesis II was positively correlated with age (Figure 2d). Our results suggest that the overall metabolic composition of microbes in BALF is not related to basic clinical information and indicates a stable state.

**Figure 2.**
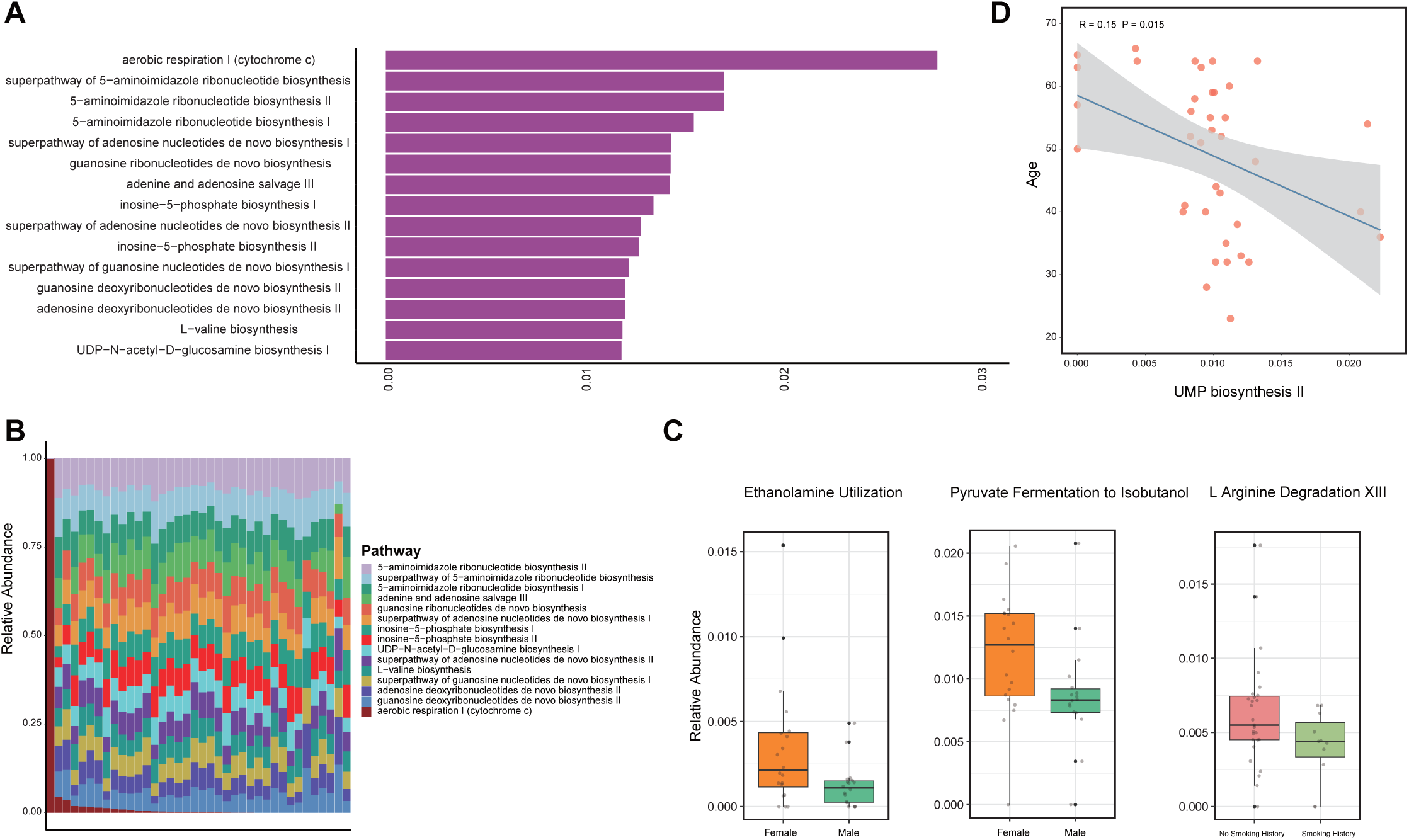
Overview of the metabolism composition of healthy individuals. A. Top 15 metabolic abundances of all samples in the cohort are sorted by the mean relative abundance of metabolism composition in each person. The scale of 0–1 corresponds to 0–100% abundance. B. Top 15 metabolic abundance compositions of all samples in the cohort. Each vertical line indicates one sample. C. Box plots illustrating variations in relative abundance for metabolic compositions between different groups. D. UMP biosynthesis II was significantly correlated with Age. The shadings indicate the 95% confidence intervals.

### Reconstruction of microbial genomes from the healthy human lung

The workflow of binning is shown in Figure 3a. Firstly, assembly was performed based on single-sample and co-sample strategies after quality control on raw sequencing data and removing human contamination. The density of N50, the number of contigs, the length of the largest contigs, and the total length were shown in Figure 3b and Figure S3a. The length of the largest contig and the total length of contigs from co-samples were larger than those from single samples. Microbial genomes representing individual bacterial species were then constructed from the assembled metagenomic sequencing data obtained from the 43 samples described above. We used the metaWRAP-Binning module to generate 844 bins from single-sample assemblies and 212 bins from mixed-sample assemblies. After dereplication, aggregation, and scoring filtering, we observed 121 bins. Furthermore, we obtained 84 metagenome-assembled genomes (MAGs) after quality assessment, meeting the criteria of >50% completeness and <10% contamination. These reconstructed microbial genomes underwent redundancy removal (average nucleotide identity ANI >0.95). A final set of 56 non-redundant MAGs were obtained. Particularly, 14 MAGs were from co-assembly (Figure S3b), indicating that co-assembly expands the number of detected MAGs. Among these 56 MAGs, 27 MAGs met the medium quality criteria (>50% completeness and <10% contamination), while 29 MAGs exhibited high quality (>90% completeness and <5% contamination). Furthermore, various indexes of high-quality MAGs were significantly higher than the medium-quality ones, including completeness, contamination, contig number, and the length of the largest contig length (Figure S3c).

**Figure 3.**
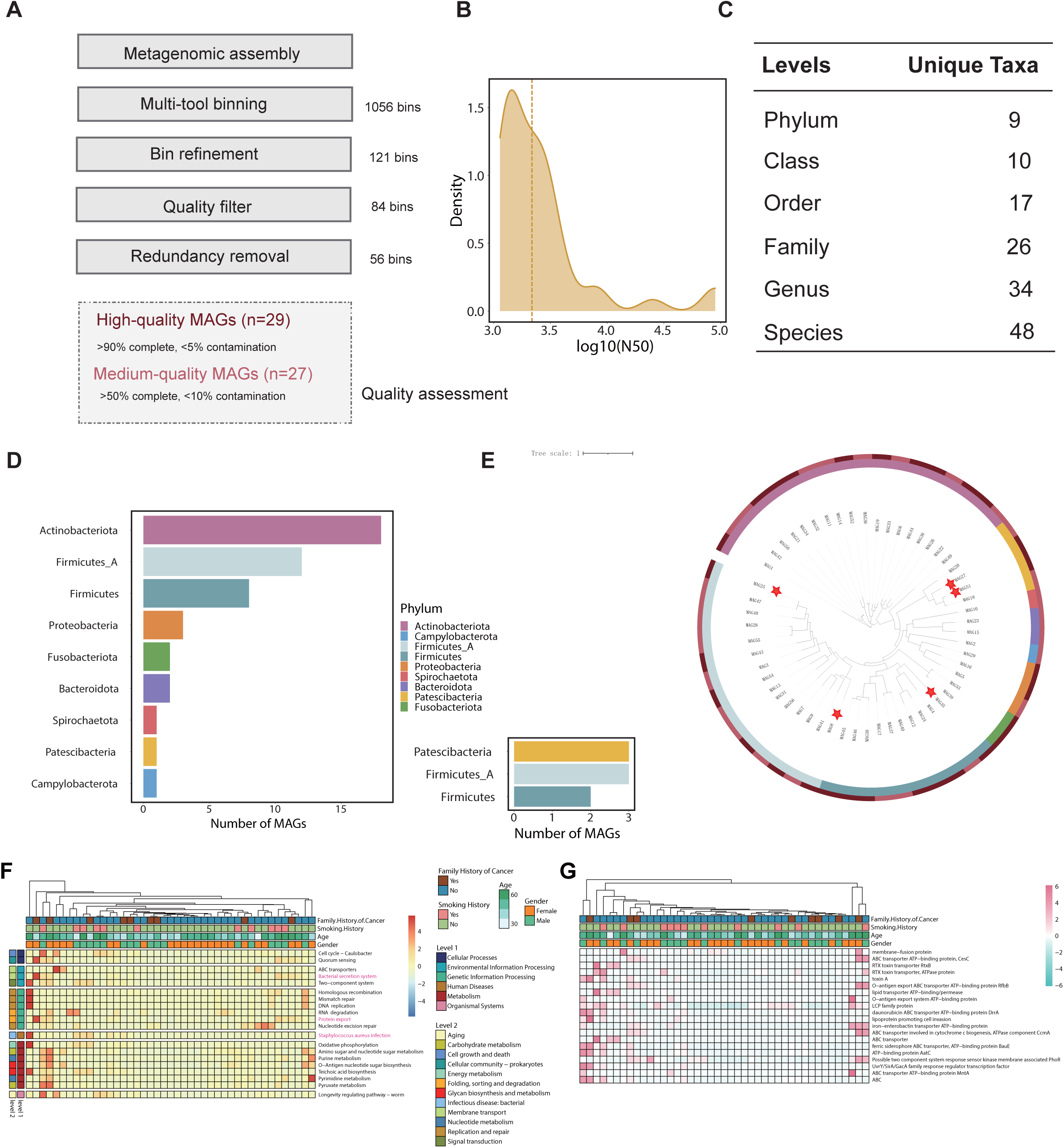
Taxonomic annotation and phylogenetic tree of 56 metagenome-assembled genomes (MAGs). A. A schematic diagram of the pipeline for constructing MAG from contigs based on the metagenomes of 43 healthy individuals. B. Statistical distribution of log-scaled N50 for single-sample assembly per sample, with dashed lines indicating the log-scaled N50 value of co-assembly. C. Taxonomic classification of 56 MAGs at various levels. D. The number of MAGs at each phylum level. E. Phylogenetic distribution of 56 MAGs. The inner-colored strips represent phyla. The outer rings indicate whether the quality of a MAG is high or medium. The star labeled MAGs represent new species. F. Heatmap of the distribution of pathways. The vertical axis represents six different kinds of pathways in level 1 and 12 different kinds of pathways in level 2. Annotation results were obtained using KEGG. G. Heatmap illustrating the distribution of virulence factors.

All MAGs exhibited a comparatively high prevalence in the 43 metagenomes (Figure S3d), indicating that these MAGs were likely strains of core species. The 56 MAGs were subsequently classified into taxa using the Genome Taxonomy Database Toolkit (GTDB-Tk) (Table S5). Further analysis showed that 48 out of the 56 MAGs were identified at the species level (Figure 3c). As shown in Figure 3d, those MAGs covered nine bacterial phyla. Most MAGs belonged to *Actinobacteriota* (18 MAGs), followed by *Firmicutes_A* (12 MAGs), *Firmicutes* (eight MAGs), *Proteobacteria* (three MAGs), *Fusobacteriota* (two MAGs), *Bacteroidota* (two MAGs), *Spirochaetota* (one MAG), *Patescibacteria* (one MAG), and *Campylobacterota* (one MAG). Among those phyla, *Actinobacteriota, Firmicutes, Proteobacteria, Fusobacteriota, Bacteroidota,* and *Spirochaetota were also detected in K*raken2, which belonged to the 196 species set. Moreover, we successfully identified *Corynebacterium argentoratense and Haemophilus_A parahaemolyticus*, which had low mean relative abundance (<1%).

### New species identified in healthy individual lung

Meanwhile, we uncovered some fascinating results regarding the identification of new bacterial species. The other eight MAGs not identified at the species level were defined as potential novel species and assigned to three phyla: *Patescibacteria* (three MAG), *Firmicutes_A* (three MAG), and *Firmicutes* (two MAG) (Figure 3d). The phylogenetic tree of the 56 MAGs was constructed based on 120 conserved proteins. The taxonomy of these MAGs at the phylum level was consistent with the phylogenetic tree (Figure 3e, Table S4). Particularly, three out of four MAGs in *Proteobacteria* were potential new species. Five out of the eight potential MAGs were highly qualified and had an ANI value less than 0.95. The five MAGs were further compared with bacterial genomes recently reported from two cohorts [14, 15]. Among these bacterial species, ANI with these five MAGs was less than 0.95, indicating that they represent unknown species identified for the first time in this study.

### Functional characterizations of high-quality MAGs

We analyzed the gene functions of 29 high-quality MAGs to gain a better understanding of the functions of the lung microbiota. At KEGG level 1, there were six identified pathways, including metabolism (109 pathways in level 2), human diseases (32 pathways in level 2), organismal systems (27 pathways in level 2), environmental information processing (13 pathways in level 2), cellular processes (12 pathways in level 2), and genetic information processing (12 pathways in level 2). In total, we identified 205 pathways at level 2. Among these, 118 pathways (57.56%) were consistently present in all samples, indicating that these pathways represent core functions of the lung microbiota. The most abundant pathways were depicted in Figure 3f. The largest number of pathways in level 2 were related to metabolism and genetic information processing, indicating their critical role in the lung micro-ecosystem. Particularly, the bacterial secretion system, protein export, and *staphylococcus aureus* infection were also detected, indicating the cross-talk between bacteria and the host. The observation of *staphylococcus aureus* infection is supported by the high relative abundance of *staphylococcus aureus* detected using Kraken2.

The prevalence of viral factors from 29 highly qualified MAGs was assessed based on the Virulence Factor Database (VFDB). A total of 575 virulence factors were identified. The top 20 high-abundance virulence factors are shown in Figure 3g, including ABC transport-associated virulence genes.

### Taxonomic and functional profiles indicate minimal alterations in COVID-19 and NSCLC

To explore the differences in microbial profiling between healthy individuals and lung-diseased patients, we selected COVID-19 and NSCLC as representatives of infectious disease and cancer, respectively. We identified 212 species from 23 COVID-19 patients, 28 NSCLC patients, and 43 individuals in the healthy group after quality control and removal of human reads. Overall, the lung microbiome compositions, based on the beta-diversity metrics of the three groups, were minimally different (Figure 4a). Moreover, COVID-19 samples were more scattered, while normal samples cluster closer together. The microbiome pairwise distances of NSCLC participants were more similar to those of the healthy group than those of individuals with COVID-19 (Figure 4b). Meanwhile, the top 15 species with high relative abundance in the healthy group changed more significantly in the COVID-19 group compared to NSCLC (Figure 4c).

**Figure 4.**
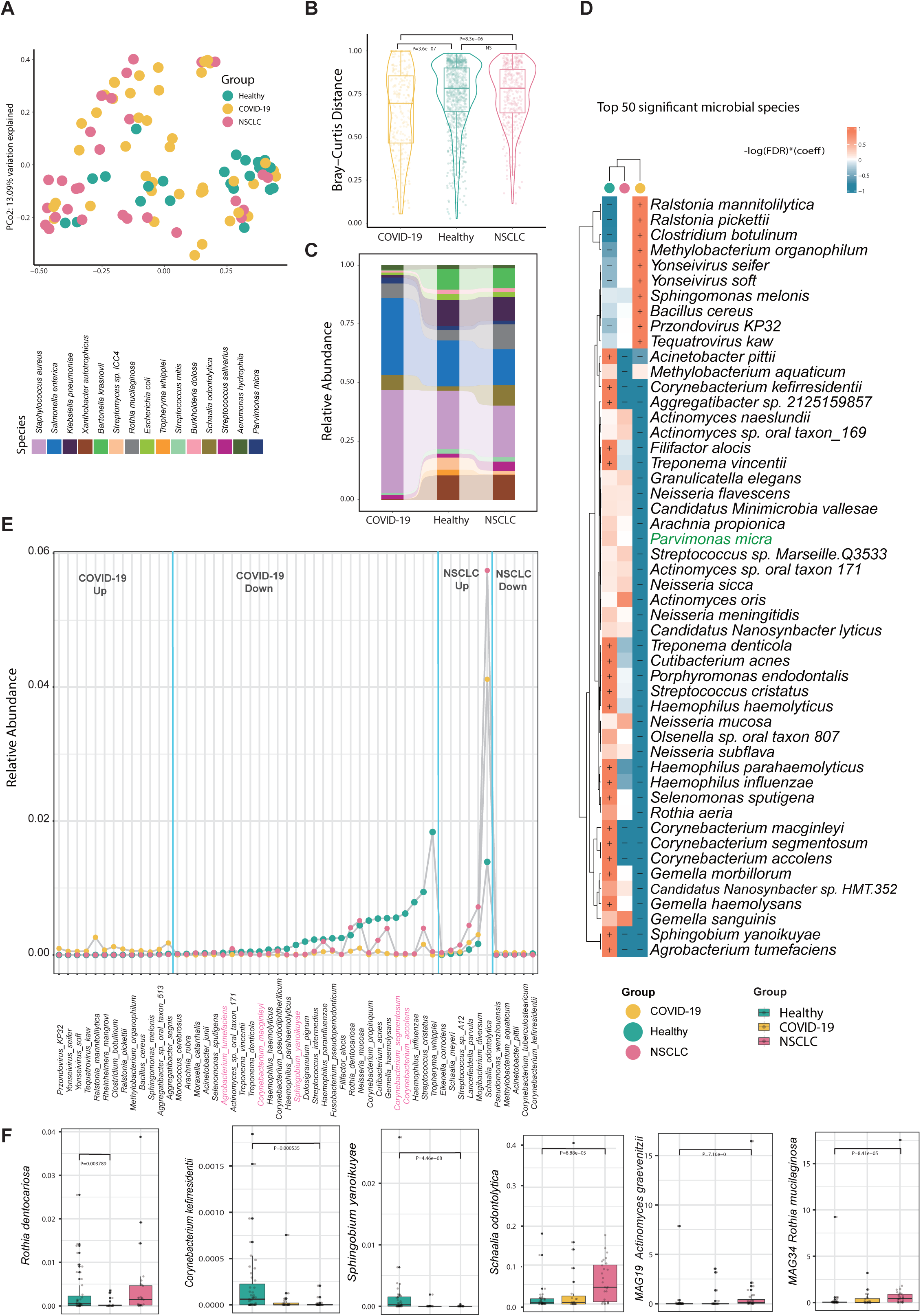
Taxonomic and functional profiles indicate changes in COVID-19 and NSCLC compared to healthy individuals. A. PCoA visualizing the beta-diversity of the three cohorts. Colors indicate different groups. B. Bray-Curtis dissimilarity comparisons of three groups. The center line represents the median, the box limits indicate the upper and lower quartiles, and the whiskers show 1.5 times the interquartile ranges. C. Histogram showing the relative abundance of the top 15 species with high relative abundance from the healthy group across three groups. D. Principal microbial species associated with health status, COVID-19, and NSCLC. The top 50 microbial species were clustered based on taxonomy. Associations were color-coded based on the direction of effect (red for positive, blue for negative) and coefficient. Significant associations at FDR < 0.05 were indicated with a plus sign (for positive correlations) or a minus sign (for negative correlations). E. Dot plots showing relative abundance of species significantly (|FoldChange| > 2 and Wilcox test P value < 0.01) enriched in COVID-19 and NSCLC. F. Box plots illustrating variations in relative abundance for species and MAGs among three groups. **Abbreviations:** PCoA, Principal coordinate analysis; NSCLC, non-small cell lung cancer.

To further explore the microbiota changes between healthy and lung-diseased individuals, MaAsLin2 analysis was conducted based on species detected via Kraken2. We identified 47 high and 14 lower species in the comparison between the healthy group and the lung-diseased group, 23 high and 118 lower species in the COVID-19 versus healthy group comparison, and 44 high and 33 lower species in the NSCLC versus healthy group comparison (Table S6). Among the top 15 high-abundance bacteria in the healthy group, *Escherichia coli*, *Tropheryma whipplei*, and *Aeromonas hydrophila* were significantly more abundant in the healthy group compared to the lung disease group. Seven out of 32 representative species in the healthy group showed enrichment compared to the lung disease group, emphasizing their potential role in maintaining lung homeostasis. The top 50 significantly different microbial species were displayed in Figure 4d. Among those species, 20 were significantly enriched in the healthy group, while 30 were in the lung disease groups. Among those 50 species, *Parvimonas micra* also belonged to the 15 high-abundance species in the healthy group.

Next, we calculated differences in the relative abundance of bacteria among the three groups with a threshold of absolute value of fold change >1 and P-value < 0.01 using the Wilcoxon test (Table S7 and S8; Figure 4e-f). *Rothia dentocariosa*, *Corynebacterium kefirresidentii, Sphingobium yanoikuyae* were enriched in the healthy group, while *Schaalia odontolytica* was enriched in the NSCLC group. According to a case report, *Rothia dentocariosa* was cultured from BALF and caused opportunistic pulmonary infection [16]. *Corynebacterium kefirresidentii* was identified as a dominant member of the human skin microbiome through 16SrRNA sequencing [17]. *Sphingobium yanoikuyae* can degrade carcinogenic products and may reduce in gastric cancer [18]. *Schaalia odontolytica* was linked to resistance to neoadjuvant chemoradiotherapy in rectal cancer [19]. The relative abundance of MAGs among the three groups was also assessed (Table S9 and S10; Figure 4f). *Actinomyces graevenitzii* and *Rothia mucilaginosa* were significantly enriched in NSCLC. *Actinomyces graevenitzii* was isolated and cultured from the BALF of a patient with pulmonary actinomycosis [2].

## Discussion

This study provides a comprehensive dataset of microbiota, an exhaustive catalog of MAGs, identifies potential new species in healthy lungs, and explores microbial alterations in lung diseases. Microbial profiling of 196 non-low-abundance species from 43 healthy individuals was conducted. The 15 dominant species consist of communal pathogens, opportunistic pathogens, and non-pathogenic commensals. For instance, *Staphylococcus aureus* is detected in 30% of the population, mostly colonized in the nasal, throat, skin, and gastrointestinal tract [20], and has the potential to cause pneumonia. *Klebsiella pneumoniae* has emerged as a major clinical and public health threat due to the rise of multidrug-resistant strains [21]. Interestingly, *Rothia mucilaginosa* has an anti-inflammatory effect induced by pathogens or lipopolysaccharides in chronic lung disease [22]. Proteins produced by *Streptococcus mitis* and *Streptococcus oralis* from viral pneumonia patients significantly increased virus replication in lung epithelial cells [23]. And *Streptococcus mitis* was linked to preserved lung function and favorable survival [24].

Therefore, our results indicated that the dominant and keystone bacterial species consisted of both pathogens and non-pathogens, suggesting a complex microbial community in the lower respiratory tract of healthy lungs. It has been reported that pathogens, including *Streptococcus pneumoniae* and *Haemophilus influenzae* were found to be colonized in the airways of 20-50% of healthy individuals [25]. With an increasing number of bioinformatic methods being utilized to distinguish pathogens from background microorganisms, our findings may assist in identifying pathogens in the clinical diagnosis of pulmonary infections[26].

We also identified a significant amount of oral-associated microbiota in the lower respiratory tract (LTR) of healthy adult participant. In keystone species obtained through co-occurrence network analysis, *Granulicatella adiacens* belongs to microbiota in the oral cavity, urogenital tract, and intestinal tract [27] and can cause periodontitis [28]. *Mogibacterium diversum* was detected in human saliva samples [29]. *Salmonella enterica* has been isolated from stool samples of individuals with diarrhea in Grace[30]. Meanwhile, the dominant species *Streptococcus mitis, Streptococcus oralis* were also oral involved. It have been reported that oral bacteria are likely the source of lung microbiota in healthy individuals [31]. It was concluded that an increased presence of oral microbes entering the lungs was associated with reduced lung function and elevated levels of pro-inflammatory cytokines [32].

Our study provided exhaustive genomics datasets for the lung microbiota. The co-assembly strategy increased the recovery ratio of MAGs, for instance, four of 29 high-quality MAGs were obtained using the co-assembly method. Furthermore, one out of five new species originated from that method, indicating that co-assembly enhanced the potential for reconstructing novel genomes. It was reported that co-assembly is an effective approach for reconstructing low-abundance MAGs [33]. In this study, MAG6, MAG8, MAG9, and MAG12 were obtained through the co-assembly method, and their abundance was lower at the individual level compared to MAGs from the single-sample assembly. Moreover, in 29 high-quality MAGs, 15 were detected by Kraken2. Six were identified at the genus level, three were not identified at the genus level, and five new species were discovered. This indicates that assembly expanded the scope of microbiota identification. Meanwhile, *Tropheryma whipplei* (MAG1), *Rothia mucilaginosa* (MAG34), and *Parvimonas micra* (MAG56), which belonged to 29 MAGs, were detected among the top 15 high-abundance species according to Kraken2, highlighting their significant role in lung microecology.

We observed that the microbiota of the diseased lung was altered compared to a healthy state, and the abundance of bacteria varied between the two different lung diseases. Interestingly, COVID-19 patients have a more distinct profile compared to NSCLS. For instance, individuals with COVID-19 had smaller Bray-Curtis distance values and experienced more microbial disturbance. Our results suggest that the microbial communities in infection-induced lung diseases may be more pronounced than in lung cancer, which requires further data for confirmation.

There are several limitations in our study. First, the sample size was not large enough to elucidate the lung micro-ecology. Second, each sample needs a blank control experiment to eliminate environmental contamination. Third, some crucial species need to be validated through experiments. Overall, we present microbial abundance and MAGs, which will significantly enhance the capacity to conduct taxonomic grouping and metagenomic analyses for future studies on the pulmonary microbiota, as well as the associations between health and diseases. We also lay the groundwork for the future development of more personalized and precise management and therapy strategies for lung health.

## Declarations

### Author statement

**Cheng Cheng** Methodology, Formal analysis, Writing – Original Draft. **Yangqian Li** Investigation, Resources, Writing – Original Draft. **Suyan Wang** Data Curation, Visualization **Haoyu Wang** Data Curation, Resources, Investigation. **Dan Liu** Supervision. **QingLan Wang** Supervision, Data Curation. **You Che** Supervision, Methodology. **Linlin Xue** Investigation. **Ningning Chao** Data Curation. **Xuan He** Supervision. **Chengping** Data Curation. **Huohua Tian** Investigation. **Jing Zhou** Investigation. **Xin Wang** Data Curation. **Yurui Yang** Data Curation. **Yuqi Zhu** Data Curation. **Renjie Xu** Data Curation. **Zhoufeng Wang** Supervision, Resources, Writing – Review & Editingm. **Weimin Li** Conceptualization, Supervision, Funding acquisition

## Acknowledgments

We are grateful to all participants who volunteered for this study.

## Data sharing statement

The raw metagenomic sequencing data generated in this study was available in the National Genomics Data Center (NGDC) Genome Sequence Archive (GSA) database (https://bigd.big.ac.cn/gsa-human/) under accession number HRA008205.

## Figure Legends

**Figure S1.**
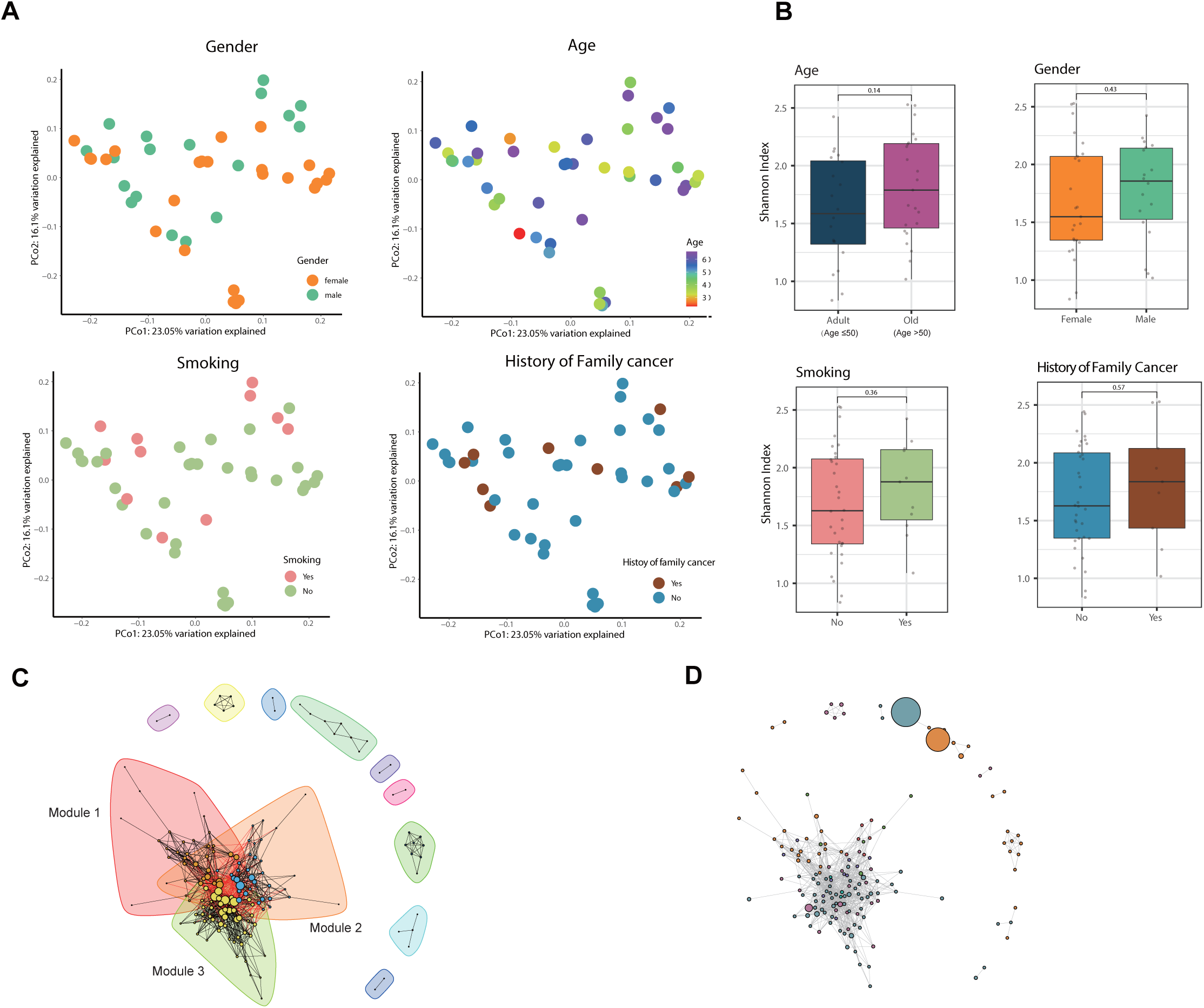
Overview of the micro-biome composition of healthy individuals, related to Figure 1. A. PCoA visualizing the beta-diversity of the 43 BALF samples from healthy volunteers grouped by gender, history of smoking, history of family-member cancer and ages. B. Alpha-diversity of individuals grouped by gender, age, smoking history, and family cancer history. P-value was calculated using a two-sided Wilcoxon test. C,D. Co-occurrence network of 155 species based on Spearman correlation coefficients among those species. Each node represents one species. Ecological relationships are represented by edges connecting two nodes. Grey lines indicate a positive correlation. In (C), Each node’s size is proportional to the rank value; The background color indicates that 12 communications have been detected. In (D), the node color represent phyla; the node size is proportional to the relative abundance of the corresponding species. **Abbreviations:** PCoA, principal coordinate analysis.

**Figure S2.**
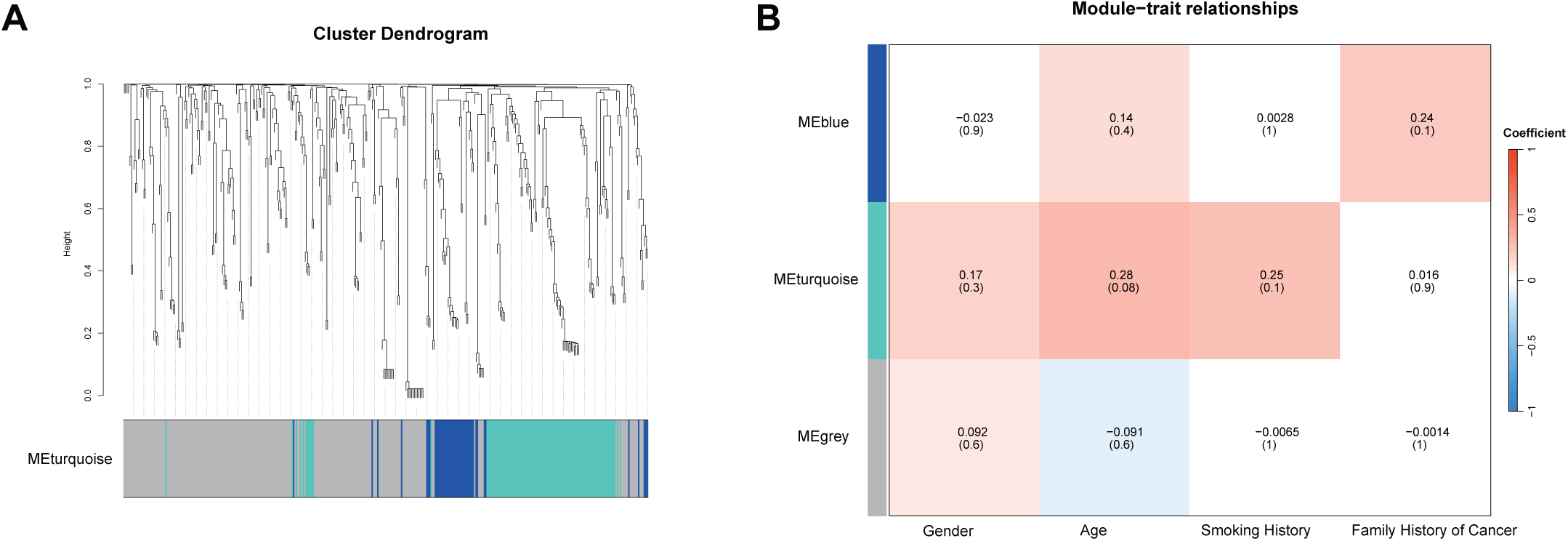
WGCNA of metabolic composition in 43 healthy individuals, related to Figure 2. A. The dendrogram and modules of metabolic pathway detected by the WGCNA. B. Pearson correlation analysis of modules and basic clinical information including age, gender, smoking history and family history of cancer. **Abbreviations:** WGCNA, weighted gene co-expression network analysis.

**Figure S3.**
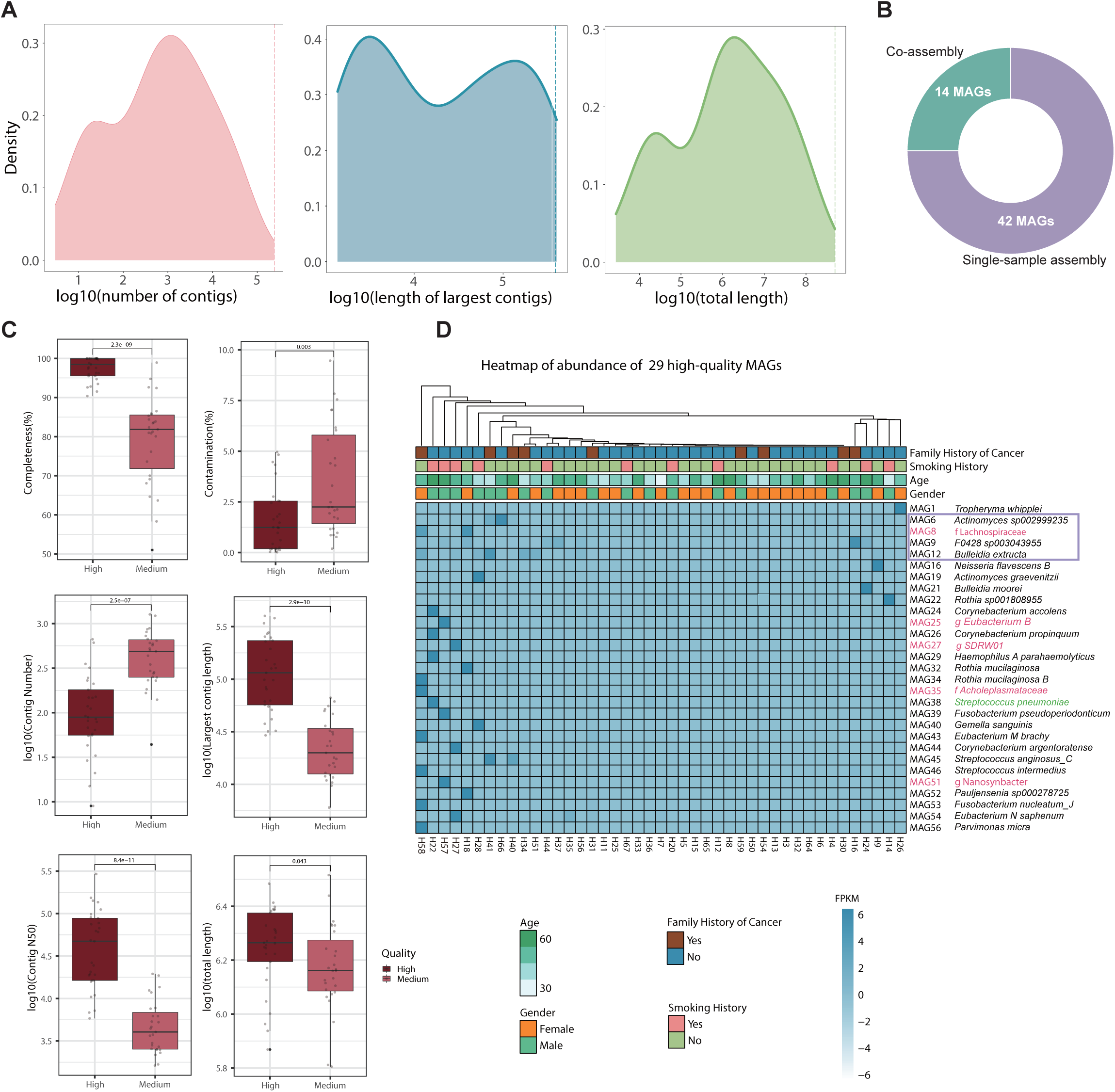
Taxonomic annotation and phylogenetic tree of 56 MAGs, related to Figure 3. A. Distribution of contig information for single-sample assembly per sample, including the number of contigs, length of the largest contig, and total length of all contigs. The dashed lines represent values of co-assembly. B. The fraction of MAGs from the single-sample assembly and co-assembly. C. Quality metrics across high-quality (n = 29) and medium-quality (n = 27) MAGs. D. Heatmap showing the abundance of 29 high-quality MAGs. MAGs in the purple frame were from co-assembly. MAGs in red indicate novel species. MAGs in green indicate species ranked 18th based on relative abundance using Kranken2. **Abbreviations: MAGs,** metagenome-assembled genome.

